# Characterization of influenza B virus variants with reduced neuraminidase inhibitor susceptibility

**DOI:** 10.1101/334201

**Authors:** R Farrukee, AE Zarebski, JM McCaw, JD Bloom, PC Reading, AC Hurt

**Affiliations:** WHO Collaborating Centre for Reference and Research on Influenza, at the Peter Doherty Institute for Infection and Immunity, Melbourne, Victoria, 3000, Australia; Department of Microbiology and Immunology, The University of Melbourne, at the Peter Doherty Institute for Infection and Immunity, Melbourne, Victoria, 3000, Australia; School of Mathematics and Statistics, The University of Melbourne, Melbourne, Australia; Centre for Epidemiology and Biostatistics, Melbourne School of Population and Global Health, The University of Melbourne, Melbourne, Australia; Victorian Infectious Diseases Reference Laboratory Epidemiology Unit, at the Peter Doherty Institute for Infection and Immunity, Melbourne, Victoria, 3000, Australia; Infection and Immunity theme, Murdoch Children’s Research Institute, Royal Children’s Hospital, Melbourne, Australia; Division of Basic Sciences and Computational Biology Program, Fred Hutchinson Cancer Research Center, Seattle, WA, USA

## Abstract

Treatment options for influenza B virus infections are limited to neuraminidase inhibitors (NAIs) which block the neuraminidase (NA) glycoprotein on the virion surface. The development of NAI resistance would therefore result in a loss of antiviral treatment options for influenza B infections. This study characterized two contemporary influenza B viruses with known resistance-conferring NA amino acid substitutions, D197N and H273Y, detected during routine surveillance. The D197N and H273Y variants were characterized *in vitro* by assessing NA enzyme activity and affinity, as well as replication in cell culture compared to NAI-sensitive wild-type viruses. *In vivo* studies were also performed in ferrets to assess the replication and transmissibility of each variant. Mathematical models were used to analyse within-host and between-host fitness of variants relative to wild-type viruses. The data revealed that the H273Y variant had similar NA enzyme function relative to its wild-type but had slightly reduced replication and transmission efficiency *in vivo*. The D197N variant had impaired NA enzyme function but there was no evidence of reduction in replication or transmission efficiency in ferrets. Our data suggest that the influenza B variant with H273Y NA substitution had a more notable reduction in fitness compared to wild-type viruses than the influenza B variant with the D197N NA substitution. Although a D197N variant is yet to become widespread, it is the most commonly detected NAI-resistant influenza B virus in surveillance studies. Our results highlight the need to carefully monitor circulating viruses for the spread of influenza B viruses with the D197N NA substitution.

## Introduction

Influenza A and B viruses co-circulate in the human population and cause yearly epidemics worldwide. Even though influenza B viruses generally cause a milder disease than influenza A viruses, in certain seasons they can contribute to a substantial burden of disease and up to 25% of influenza-related mortality (1, 2). Vaccines are the main control measure for influenza disease prevention, but a recent analysis of data between 2004-2014 revealed that the effectiveness of the vaccine against influenza B infections was only 54% (3).

The neuraminidase inhibitors (NAIs) (zanamivir, oseltamivir, laninamivir and peramivir) are a class of antiviral drugs that target the conserved amino acid residues of the viral neuraminidase (NA) active site and competitively inhibit enzyme function (4, 5). However, mutations in the NA enzyme active site or adjacent framework residues can abrogate NAI interaction with the NA and reduce the susceptibility of viruses to one or more of these drugs (6). Antiviral options for the treatment of influenza B viruses are limited to NAIs as the older class of influenza antivirals, the adamantanes, are only effective against influenza A viruses (6). Development of NAI resistance in influenza B viruses is therefore a public health concern as it would remove a valuable treatment option.

*In vitro* studies consistently demonstrate that compared to influenza A viruses, influenza B viruses exhibit reduced sensitivity to oseltamivir, suggesting that the drug may already have reduced effectiveness against influenza B viruses (6-10). The clinical relevance of this has not been fully elucidated, but in 7 out of 9 clinical studies it was shown that oseltamivir treatment resolved symptoms faster in influenza A patients than in influenza B patients (11). Considering this, it is possible that NA mutations that only moderately alter the oseltamivir susceptibility of influenza B viruses may have a significant impact on the clinical effectiveness of the drug.

A number of different NA substitutions at conserved amino acid positions (e.g. E117, D197, I221 and H273) have previously been described to confer reduced inhibition by the NAIs *in vitro* (8, 12-21), but the impact of these substitutions on enzyme function, virus replication or transmissibility, has only been assessed in a limited number of studies (14, 22, 23). The fitness of influenza B viruses with either the H273Y or D197N NA substitution is of particular interest as a number of viruses with either substitution have been recently found in patients in community settings who, unlike hospitalized or immunocompromised patients, do not typically receive NAI treatment (8, 9, 17, 18, 24). Two reports have identified household transmission of influenza B viruses with the D197N NA substitution (18, 25) and more recently, a global surveillance report identified a cluster of six influenza B viruses with the D197N NA substitution in Japan in early 2014, further suggesting potential community transmission of the variant virus (18). Interestingly, 22 out of the 27 viruses with the D197N substitution reported in the literature were from the B/Yamagata-lineage (17, 18, 25-30). There has also been examples of suspected transmission of influenza B viruses with the H273Y NA substitution (9). The H273Y NA substitution in influenza B viruses occurs at the equivalent residue to that of the H275Y NA substitution in A(H1N1) viruses, which was present in the oseltamivir resistant A(H1N1) viruses that spread globally in 2008/9 (31, 32).

The effect of H273Y NA substitutions in influenza B viruses has been previously studied using reverse genetics (rg) in the B/Yamanashi/166/98 virus background (15, 22, 23). To date, few studies have reported the effect of the H273Y or the D197N NA substitution in contemporary viruses, which is important because it has been shown that the fitness consequences of resistance-conferring mutations can vary due to the genetic background of the NA (33, 34). Although experiments using reverse genetics can be useful in defining the impact of a single mutation on viral fitness; they don’t evaluate the effect of the rest of the viral genome that may play an important role in the fitness of that virus. Our aim was to characterize two ‘naturally’ occurring influenza B variant viruses containing either the H273Y or D197N NA substitutions which had been detected during routine surveillance in patients not being treated with NAIs, compared to closely matched wild-type viruses by assessing their enzyme function, *in vitro* replication and *in vivo* replication and transmission.

## Results

### NAI susceptibility, NA activity, surface expression and substrate affinity

The effect of the D197N and H273Y substitutions on NA enzyme function were assessed using four different assays. The MUT-Y273 variant had a 3-fold increase in oseltamivir IC_50_ and an 85-fold increase in peramivir IC_50_ compared to WT-H273, but the IC_50_ for zanamivir and laninamivir were not significantly different (Table I). The MUT-Y273 virus had comparable K_m_ (substrate affinity) to that of the WT-H273 virus (Table I). Similarly, the relative NA surface expression and enzyme activity of the MUT-Y273 virus compared to the WT-H273 virus was 115 ± 13.4% (mean ± SEM) and 119 ± 23.1% respectively, neither of which was significantly different (Figure I).

**Figure I:**
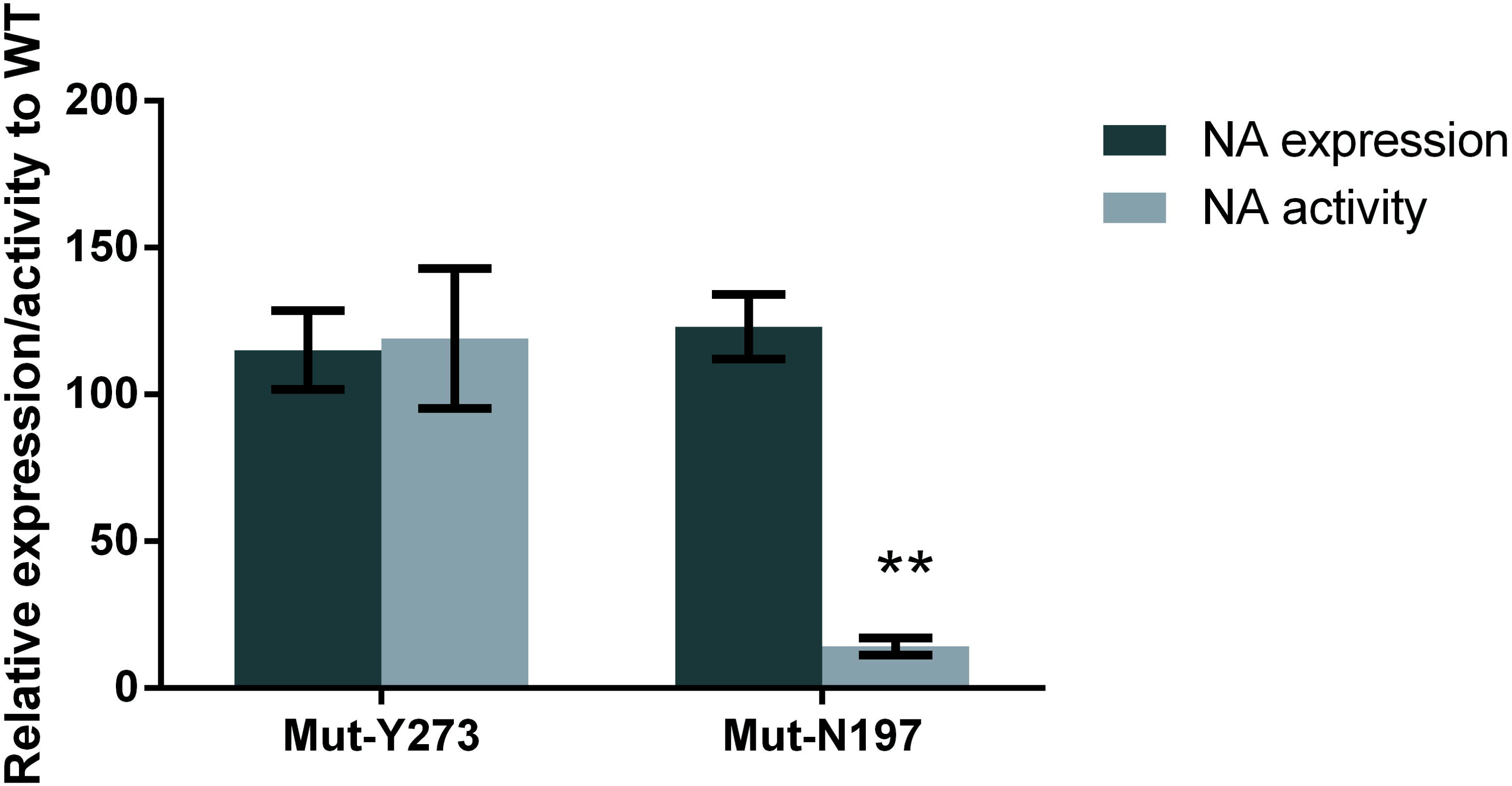
The mean NA surface expression and activity of influenza B variants relative to the corresponding WT. HEK-293T cells were transfected with plasmids containing the NA gene encoding for WT and variant proteins. At 20 hours post-transfection cells were analysed for NA activity using a MUNANA-based assay, and NA expression using a FACs-based assay. The activity and expression data of the variants were normalized relative to their corresponding WT and expressed as the mean±SEM. Data are derived from two independent experiments, each containing triplicate samples and compared between relevant groups using the non-parametric Mann-Whitney’s U-Test.

**Table I.**
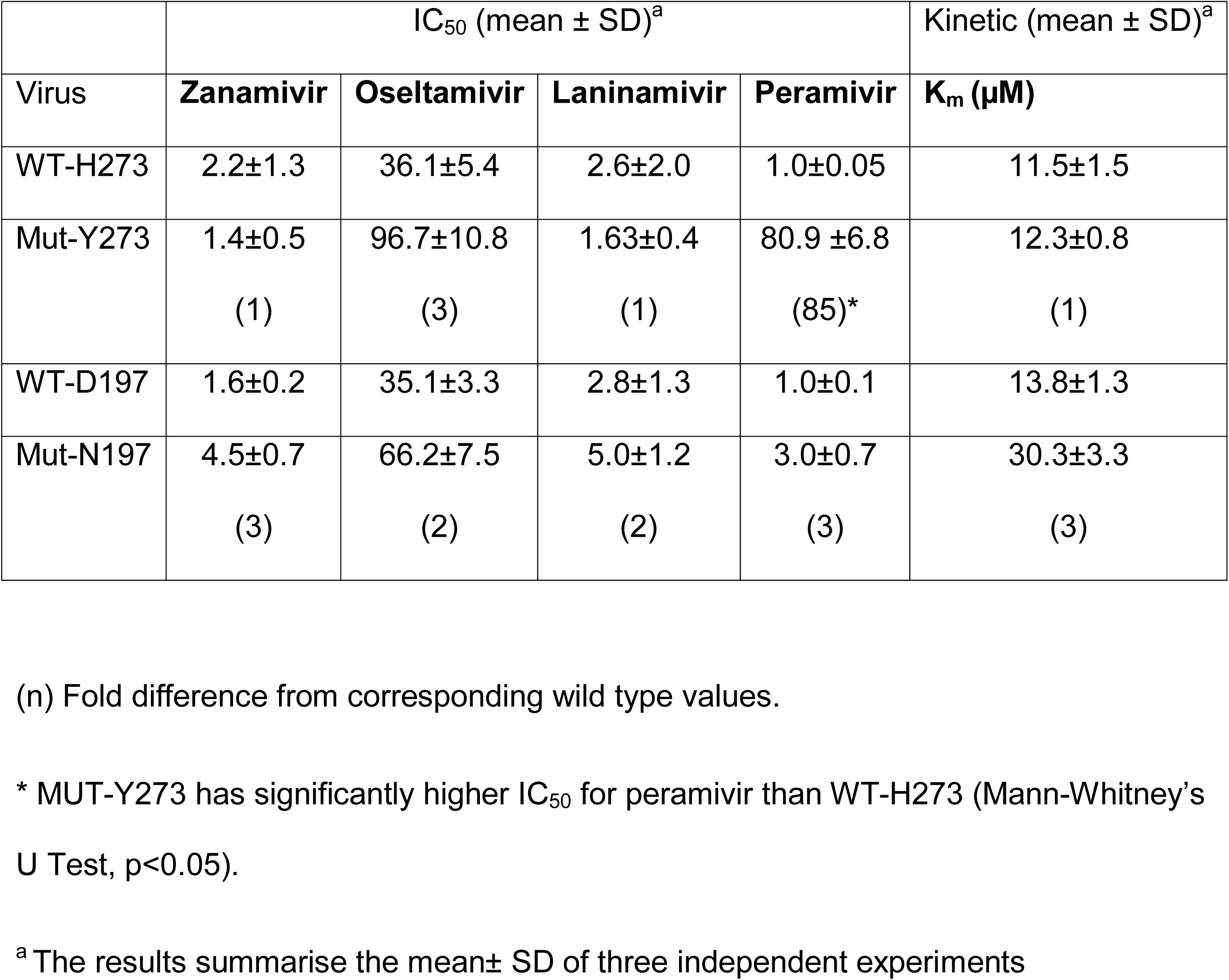
Effect of neuraminidase mutation on IC_50_ value and enzyme kinetics

|  | IC<sub>50</sub> (mean ± SD)<sup>a</sup> |  |  |  | Kinetic (mean ± SD)<sup>a</sup> |
| --- | --- | --- | --- | --- | --- |
| Virus | Zanamivir | Oseltamivir | Laninamivir | Peramivir | K<sub>m</sub> (μM) |
| WT-H273 | 2.2±1.3 | 36.1±5.4 | 2.6±2.0 | 1.0±0.05 | 11.5±1.5 |
| Mut-Y273 | 1.4±0.5<br>(1) | 96.7±10.8<br>(3) | 1.63±0.4<br>(1) | 80.9 ±6.8<br>(85)* | 12.3±0.8<br>(1) |
| WT-D197 | 1.6±0.2 | 35.1±3.3 | 2.8±1.3 | 1.0±0.1 | 13.8±1.3 |
| Mut-N197 | 4.5±0.7<br>(3) | 66.2±7.5<br>(2) | 5.0±1.2<br>(2) | 3.0±0.7<br>(3) | 30.3±3.3<br>(3) |

(n) Fold difference from corresponding wild type values.
\* MUT-Y273 has significantly higher IC<sub>50</sub> for peramivir than WT-H273 (Mann-Whitney's U Test, p<0.05).
<sup>a</sup> The results summarise the mean± SD of three independent experiments

The MUT-N197 virus had a 2-3-fold reduction in susceptibility to all four NAIs compared to that of the WT-D197 virus (Table I). The substrate affinity (K_m_) was 3-fold lower for the MUT-N197 virus compared to the WT-D197 virus (Km values are inversely proportional to substrate affinity) but this difference was not statistically significant (Table I). The cell-surface expression of the NA enzyme of the Mut-N197 variant was 123 ± 10.9% relative to the WT-D197 virus but the NA enzyme activity of the Mut-N197 variant was only 14.2 ± 3.0%, which was significantly lower (p<0.01) than the WT-D197 virus (Figure I).

### In vitro replication kinetics

Multi-step growth curves were performed in MDCK and MDCK-SIAT (SIAT) cells to assess the *in vitro* replication kinetics of the WT and MUT viruses. The MUT-Y273 variant showed significantly reduced viral titres compared to the WT-H273 virus at 24 and 36 hours post-infection in MDCK cells and at 36 hours post-infection in SIAT cells (Figure II a, b). However, at 48 hours post-infection there were no significant differences in viral titres between the MUT-Y273 and WT-H273 viruses in either cell line.

**Figure II:**
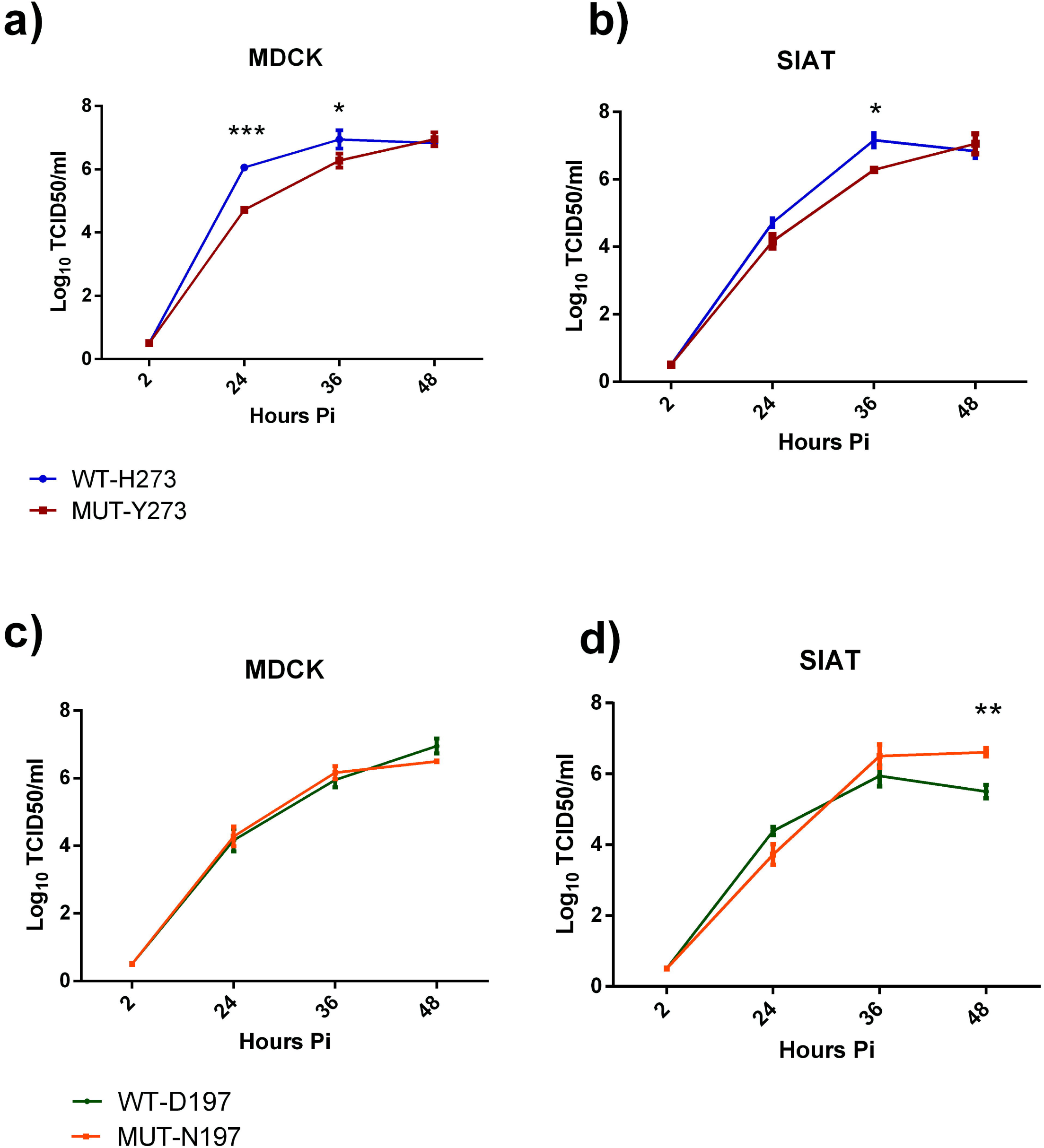
Multi-cycle replication of influenza B WT and MUT viruses in MDCK and SIAT cells. Cells were infected at a MOI of 0.01 and supernatants collected and assayed for infectious virus at 2, 24, 36 and 48 hr post-infection. Data show the mean (± SEM) of triplicate samples for Mut-Y273 vs WT-H273 (a, b) and Mut-N197 vs WT-D197(c, d). A two-way ANOVA analysis with Bonferroni’s post-test analysis was done to compare mean viral titres at each time point between WT and MUT viruses. * p<0.05, **p<0.01, ***p<0.001

The MUT-N197 variant demonstrated no reduction in replication compared to the WT virus in MDCK cells and actually outgrew the WT-D197 virus in SIAT cells at 48 hours post-infection where it reached a significantly higher titre of 6.61 ± 0.11 vs 5.5 ± 0.2 Log_10_ TCID_50_/ml (p<0.01) (Figure II c, d).

### Replication and transmission of pure viral populations in ferrets

To gain insight regarding virus replication *in vivo*, we determined titres of infectious virus in nasal wash samples taken at various times after intranasal infection of ferrets. Figure III a & c summarizes the results for animals infected with either the WT-H273 or the Mut-Y273 virus. In both donor groups (infected by direct intranasal inoculation), infectious virus was detectable until day 8 post-infection and peak viral titres were not significantly different. There was also no significant difference in the area under the curve (AUC) of TCID_50_ plots over the 10 day period (Figure III a, c). Body temperature and weight were also assessed daily after infection and no significant differences were noted between donor animals infected with WT-H273 or Mut-Y273, although there was a trend towards greater weight loss in donors infected with WT-H273 (Figure S2).

**Figure III:**
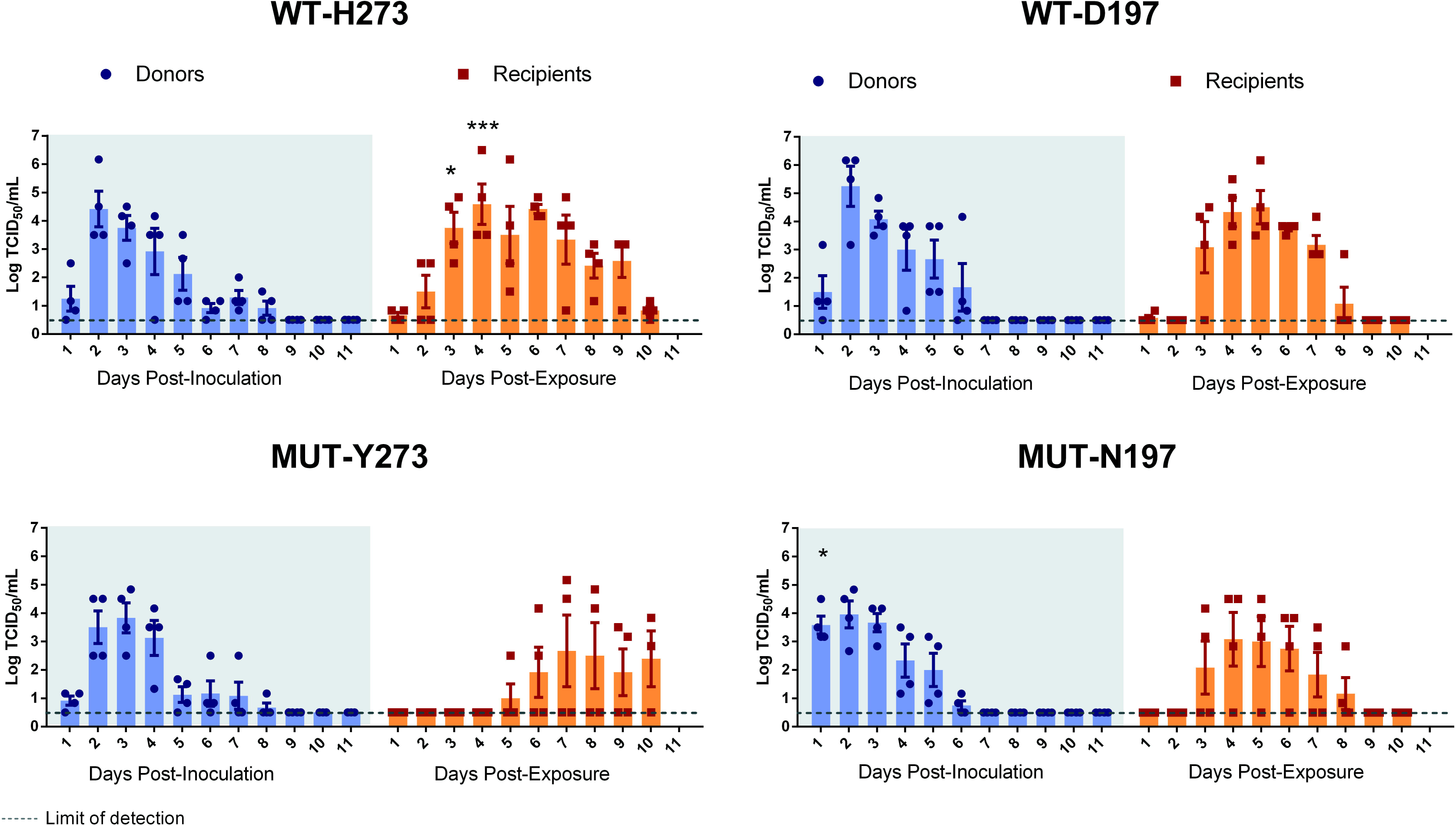
Viral shedding in nasal wash samples from donor and recipient ferrets infected with WT or MUT influenza B viruses. Donor ferrets (n= 4 per group) were experimentally infected with 10^5^ TCID_50_ of ‘pure’ WT or MUT viruses. Naïve recipients were co-housed with each donor one day after experimental infection of donors. Daily nasal wash samples were collected from both donors and recipients over a period of 11 days and titres of infectious virus in nasal wash samples were determined by TCID_50_ assay. A-D summarizes the TCID_50_ results with the bar graphs showing the mean (± SEM) titres at each day for each group. At each time point, mean viral titres were compared between donors infected with either WT or MUT virus by two-way ANOVA with Bonferroni’s post-test analysis. Similarly, analysis was done to compare titres between recipients infected with either WT or MUT virus. Significant differences in mean viral titres were observed between WT-H273 and Mut-Y273 infected recipients on day 3 and 4 post-exposure and between WT-D197 and Mut-N197 infected donors on day 1 post-inoculation. Area under the curve (AUC) of TCID_50_ graphs were also calculated and compared by Mann-Whitney’s U test; no significant differences were detected between donor animals infected with WT and corresponding MUT viruses. * p<0.05, **p<0.01, ***p<0.001.

We examined virus replication in nasal washes from naïve recipient ferrets who were co-housed with experimentally infected donors to assess the transmissibility of WT-H273 and Mut-Y273 viruses. All four recipients co-housed with WT-H273-infected donors shed infectious virus but only two out of four recipients co-housed with MUT-Y273-infected donors shed detectable infectious virus (Figure III a, c). There was a delay in transmission of the MUT-Y273 virus compared to the WT-273 virus, as peak virus titres in WT-H273-infected recipients was detected on day 4 post-exposure while the two recipients infected with the MUT-Y273 virus reached peak viral titres on day 7 post-exposure (Figure III a, c).

Next, we examined virus replication following experimental infection of animals with WT-D197 and Mut-N197 viruses. The kinetics of viral shedding was similar between donor ferrets experimentally infected with either WT-D197 or MUT-N197 viruses (Figure III b, d). The AUC of the viral replication plots and the peak viral titres between the donor groups were not significantly different. Infectious viral titres were detectable in nasal washes of at least one ferret until day 6 post-infection and donors did not experience any significant changes in either body weight or temperature (Figure S2).

The WT-D197 virus transmitted successfully to all four recipients, while the MUT-N197 virus transmitted to three out of the four recipients. However, length of viral shedding and the peak viral titres were not significantly different between recipients infected with either the WT-D197 virus or the MUT-N197 virus (Figure III b, d); there was also no transmission delay observed as with MUT-Y273. Similar to the donor ferrets, the recipient ferrets infected with either WT-D197 or MUT-N197 virus experienced no significant changes in either body weight or temperature.

Pyrosequencing analysis of relevant NA substitutions in nasal wash samples from all infected animals confirmed that neither the D197N nor the H273Y substitution was lost following replication within the airways of donor ferrets, or following transmission to recipient animals. Next generation sequence analysis on nasal wash samples from the last day of shedding in recipient ferrets was performed to confirm the genetic stability of the rest of the viral genome upon transmission. In addition to a number of synonymous mutations found across the influenza genome, the MUT-Y273 virus from one recipient ferret also contained a M403V NA amino acid substitution, and the WT-D197 virus from all recipients contained a L274I amino acid substitution in the PA. M403V in influenza B NA is distant from the enzyme active site and has only occurred on rare occasions in influenza B viruses (19 out of 5556 sequences in GISAID), while the PA substitution L274I has not been previously observed in any of the influenza B PA gene sequences on GISAID.

### Relative fitness between viral pairs based on ‘competitive mixtures’ analyses

To gain further insight into the fitness differential between each variant virus and their corresponding wild-type viruses, a competitive mixtures experiment was performed in ferrets. Mathematical models for the progression of the infection and its transmission were fitted to measurements collected during the experiment. The parameter estimates from this fitting process provide information about the relative fitness of the variant for replication within-host and its ability to transmit between hosts.

These inferences are summarized in Table II and visualizations are shown in Figure IV.

**Figure IV:**
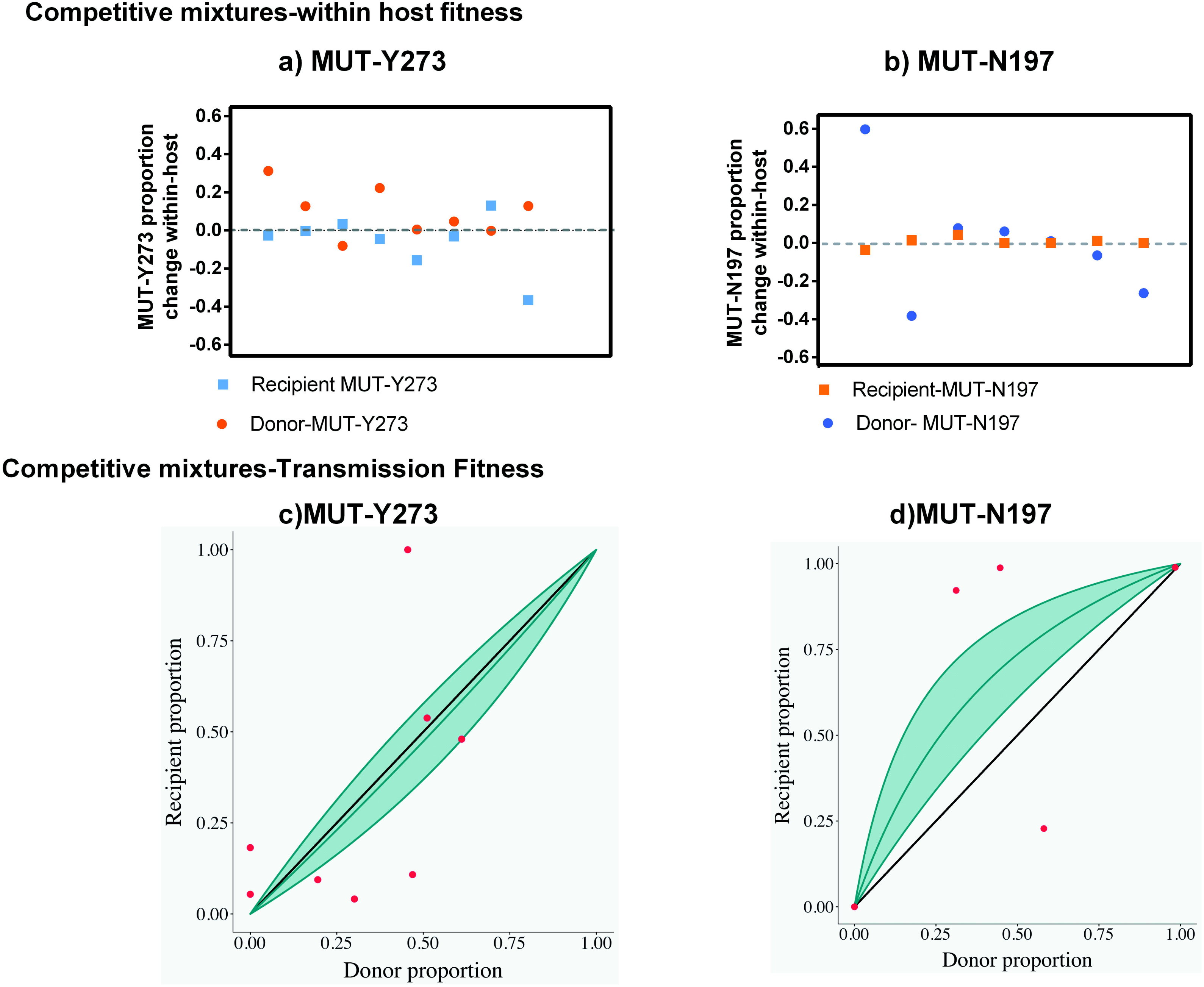
Summary of changes in mutant virus proportions in nasal washes within-host and during transmission in ‘competitive mixtures’ ferret experiments. To assess within-host and transmission fitness of mutant viruses relative to their corresponding wild-types, ferrets were experimentally co-infected with viral mixtures of WT:MUT at different ratios (50:50, 80:20 and 20:80) and co-housed with naïve recipients. The proportion of WT vs MUT was determined in the collected nasal wash samples by pyrosequencing and data were analysed only from animals where the proportion of WT and MUT could be accurately determined. A-B plots individual points that correspond to the change in the proportion of mutant virus in one ferret between the first day of infection and last day of infection (y-axis). This allows for a visual representation of within-host fitness. C-D illustrate individual points that corresponds to a transmission event between two ferrets. In these plots the abscissa is the proportion of MUT virus in donor ferrets the day preceding confirmed transmission and the ordinate shows the proportion of MUT virus in recipient ferrets within the first 24 hours of confirmed infection. The solid grey line is the representative curve of the shape parameter, calculated by fitting our data to a model described in more detail in the methods and supplementary. The 95% confidence interval is represented by the shaded region. If our fitted model lies above the symmetry line (y=x solid line) then the mutant virus is fitter than the wild-type virus, whereas if the fitted model lies below the symmetry line the converse is true.

**Table II.** Relative Within-Host and Transmission fitness between viral pairs tested in ‘competitive mixtures’ during ferret infection

|  |  | Relative within-host fitness | Relative Transmission fitness |
| --- | --- | --- | --- |
| Mutant | Wild Type | $R_0$ ratio (95% CI)<sup>a</sup> | Relative fitness value (95% CI)<sup>b</sup> |
| Mut-Y273 | WT-H273 | 0.84 (0.70,1.02) | 0.90 (0.60, 1.35) |
| Mut-N197 | WT-D197 | 0.94 (0.81,1.10) | 2.78 (1.56, 5.53) |

<sup>a</sup>If the $R_0$ ratio <1 it indicates fitness advantage of wild-type over mutant, and if $R_0$ ratio>1 the mutant is fitter than wild-type
<sup>b</sup>Relative transmission fitness value<1 indicates fitness advantage of wild type over mutant and fitness value>1 indicate mutant is fitter than wild-type

We define the relative within-host fitness as the ratio of the basic reproduction numbers of the variant and its wild-type counterpart. For the MUT-Y273 virus the Ro ratio is estimated to be 0.84 with a 2-sided 95% CI of (0.70, 1.02), with our analysis indicating that there is a 95% chance that this ratio is less than 1. So while the MUT-Y273 variant is almost certainly less fit than its wild-type counterpart, the difference is very small. There was little evidence that the mutation affected transmission fitness, with the mutant’s ability to transmit reduced by a factor of 0.90 (0.60, 1.35).

For MUT-D197, we find little evidence for a difference in within-host fitness, with an estimated ratio of 0.94 (0.81, 1.10) and a 75% chance that the variant is less fit than the wild-type. Interestingly, the transmission fitness of the MUT-N197 virus was higher by a factor of 2.78 (1.56, 5.59) over the wild-type. However, it is important to note that this estimate is driven heavily by two recipient ferrets where a large increase in the population of MUT-N197 was observed following transmission from donors infected with a 50:50 ratio of WT-D197: MUT-N197 viruses. Although the statistical model is designed to account for the variation among ferrets, we are careful to not draw too strong a conclusion on this result given the limited number of animals in the study.

A large increase in the proportion of MUT virus (>95% of population) was observed following transmission to three recipient ferrets (one in the MUT-Y273 group and two in the MUT-N197 group). To determine if non-synonymous genetic changes had been acquired across the genome of these viruses the samples were analysed by next generation sequencing. The MUT-Y273 virus in the recipient ferret had acquired a rare T90I HA substitution (contained in only 7 out of 7373 strains in GISAID), as well as the NP, PA and PB1 genes from the WT-H273 virus, which differed by 4 amino acids compared to the MUT-Y273 virus (Figure S1). In one ferret, a minor proportion (<20%) of the MUT-N197 virus population contained the NS and PA gene of WT-D197 virus (which differed by 2 amino acids compared to the MUT-N197), but the dominant MUT-N197 viral population from this ferret and from the other did not acquire any amino acid changes following transmission.

## Discussion

This study aimed to characterize two contemporary influenza B variants with either the D197N or H273Y NA substitution. The data showed that the MUT-N197 variant had significantly reduced NA enzyme activity but the replication and transmission efficiency of this virus in an *in vitro* or *in vivo* model was not notably reduced compared to a closely matched wild type virus. Further to this, mathematical analysis from competitive mixtures experiments, found little evidence of reduced relative within-host fitness and no evidence of reduced between-host transmission of the MUT-N197 virus. A previous study using an influenza B virus (B/Rochester/02/2001) with the D197N NA substitution also showed that when ferrets were infected with a mixture of wild type and variant virus, a mixture was still maintained 5-days post infection (27).

It is interesting that the significantly reduced NA enzyme activity of the MUT-N197 virus did not predict the outcome of the replication or transmission fitness of this variant in the ferret model. This may be because the enzyme-based assay is inherently limited in its ability to capture the complex interplay between the different viral proteins during replication and transmission in a biological setting. This kind of discrepancy between *in vitro* and *in vivo* data has been described previously (35), and highlights the need for caution when assessing viral fitness using only *in vitro* parameters.

As has been seen in other studies, the MUT-Y273 virus had reduced susceptibility to oseltamivir and peramivir, but not to zanamivir and laninamivir (36, 37). While the impact on peramivir binding appears greater, due to the 85-fold increase in IC_50_ (as opposed to 3-fold for oseltamivir), this is a result of substantially higher oseltamivir IC_50_ levels (∼36 nM) compared with peramivir IC_50_ levels (∼1 nM) for wildtype influenza B viruses. The actual peramivir and oseltamivir IC_50_ values for MUT-Y273 are 80.9 nM and 96.7nM respectively, showing that the *in vitro* inhibitory concentration of both drugs against the MUT-Y273 virus is similar. We show that the MUT-Y273 virus had equivalent NA activity and expression to the WT-H273 virus, but delayed growth in cell culture. Previous experiments using a rg-H273Y virus showed that relative to the rg-WT, the rg-H273Y virus had significantly higher NA activity and surface expression, and superior replication kinetics in a competitive mixtures experiment in normal human bronchial epithelial cells (14, 22, 23). The discrepancy in these fitness outcomes between our 2015 H273Y virus and a 1998 rg-H273Y virus may be due to differences in viral background (16 amino acid differences). The MUT-Y273 virus had similar replicative fitness in donor ferrets following intranasal infection viruses and delayed and reduced transmission between co-housed ferrets compared to WT-H273 virus when tested in a pure population. Genetic analysis showed that the MUT-Y273 virus which transmitted to one of the recipient ferrets contained a M403V NA change, while all of the WT-H273 viruses that infected recipient ferrets contained the L274I PA substitution. Neither of these mutations has been previously reported in the literature to alter viral fitness and therefore their role in compensating for the NA substation is unknown. Mathematical analysis showed evidence of a slight reduction in the within-host fitness of the variant in a competitive mixture with the wild-type but no evidence of reduction in between-host transmission.

There is suggestive evidence that transmission efficiency of the MUT-Y273 variant was lower when tested as a pure population compared to when assessed in a competitive mixtures experiment. It is possible that this difference is due to the reassortment with WT-H273 virus that occurred during the competitive mixtures experiments which may have led to an increase in the transmissibility of the MUT-Y273 virus. It is a limitation of this study that while the MUT viruses were closely matched to their paired WT viruses, with identical HA, NA (except for the resistance mutation of interest) and MP sequences, they did have a small number of amino acid differences in the internal genes (Figure S1). However these variant viruses represent real circulating strains and therefore have greater relevance than performing experiments using reverse genetics with laboratory reference strains.

It is important to note that our study contributes to the very limited number of animal studies that have reported influenza B virus transmission to date. One previous study demonstrated successful influenza B transmission between guinea pigs in contact and non-contact models (38), while only two studies (with mixed results) have previously reported on influenza B transmission in the ferret model (39, 40). Of these two, one study reported successful aerosol and contact transmission of a mouse-adapted influenza B virus, but could not demonstrate transmission by either model using a non-mouse adapted isolate (39). The second study using a modified influenza B virus lacking the NB protein showed that one out of the four recipient ferrets were successfully infected by aerosol transmission (40). Our results show successful contact transmission of both the influenza B wild-type viruses tested (4 out of 4 ferrets) and transmission of influenza B variants in the ferret model with circulating human influenza viruses. Given the successful results of our contact ferret transmission model using wildtype and variant strains, future evaluations of the fitness of influenza B strains with reduced NAI susceptibility should consider the use of an aerosol transmission model.

## Conclusion

The results from our experiments suggest that of two influenza B viruses with reduced NAI susceptibility, the viral fitness of a recent influenza B virus with the H273Y NA substitution was reduced compared to an equivalent wild-type virus, but an influenza B virus with a D197N NA substitution showed little reduction in reproduction or transmission in the ferret model. Although the frequency of influenza B viruses with the D197N NA substitution circulating in the community is still less than 1%, it remained the most commonly reported NAI resistant strains in recent years. Future studies to determine if the MUT-D197 virus has reduced capacity to transmit via aerosol transmission will be important.

In the future it is possible that the NA variants examined in our study may gain additional ‘permissive’ mutations that improve their fitness, as was observed with the H275Y NA substitution in seasonal H1N1 viruses (33). This may be more likely for viruses with the D197N NA substitution, which appear to have a smaller fitness deficit than viruses with the H273Y NA substitution. This knowledge, alongside the fact that clinical reports suggests that oseltamivir is less effective against influenza B and therefore minor changes in the viral NA may further reduce oseltamivir effectiveness, emphasizes the importance of continued monitoring of influenza B viruses for NA substitutions in the future.

## Materials and Methods

### Virus selection and stocks

A B/Yamagata/16/88-like Influenza B viruses with the D197N NA substitution designated B/Singapore/GP702/2015 (MUT-N197), was isolated from a 8-year old female patient not receiving NAI therapy and submitted to the WHO Collaborating Centre for Reference and Research on Influenza, Melbourne, Australia by the Ministry of Health, Singapore. Another B/Yamagata/16/88-like virus with the H273Y NA substitution designated B/Perth/136/2015 (MUT-Y273) was isolated from a 61-year old female patient not receiving NAI therapy and submitted during routine surveillance. Wild-type viruses B/Minnesota/23/2015 (WT-D197) and B/England/598/2014 (WT-H273) were kindly provided by the Centers for Disease Control and Prevention (CDC), Atlanta, United States and the Crick Institute, London, England, respectively, and were specifically chosen as wild-type pairs for MUT-N197 and MUT-Y273 respectively, due to their close genetic homology to the two variant viruses. The WT-D197: MUT-N197 pair had identical gene sequences except for the desired D197N mutation the NA, one amino acid difference in the NS gene and one amino acid difference in the PA gene Similarly the WT-H273: MUT-Y273 virus had two amino acid changes in the NP gene one change in the PB1 gene and one change in the PA gene (see Figure S1 for details). Viruses were propagated in Madin-Darby canine kidney (MDCK) cells and infectivity titres were determined by calculating the 50% tissue culture infectious dose (TCID_50_/mL) as previously described (41).

### In vitro NA inhibition and NA kinetics assay

To determine the IC_50_ (50% inhibitory concentration) of the influenza B viruses for each NAI, a fluorometric MUNANA (2-(4-methylumbelliferyl)-a-d-N-acetylneuraminic acid)-based assay was performed as previously described (42). NAI compounds were purchased from Carbosynth, UK. The Michaelis-Menten constant K_m_ (substrate affinity was also determined by measuring the rate of reaction at different MUNANA concentrations, as previously described (43), but using a MUNANA concentration range from 0-800 µM. The IC_50_ and enzyme kinetics assays were determined by three independent experiments.

### Neuraminidase surface expression and activity assay

Measurement of cell-surface NA expression and activity was performed by transfecting HEK-293T (293T) cells with an expression plasmid bearing the NA gene of interest with a C-terminal V5 epitope tag (33, 44, 45).

At 20 hours post-transfection, the NA enzyme activity on the cell surface was measured using a modified MUNANA-based assay as previously described (44). The enzyme expression in these cells was also measured by staining with an Anti-V5 Alexaflour 647 Antibody (Thermofisher, Australia) and measuring staining intensity using flow cytometry (44). The NA activity results were normalized for total surface expression and results for each variant were normalized against fluorescent staining intensity/activity of corresponding wild-type viruses. Two independent experiments were done to assess NA expression and activity, where each virus was tested in triplicates.

### In vitro replication kinetics

A multi-step replication experiment was carried out in MDCK and MDCK-SIAT1(SIAT) cell lines, which were grown to confluence in 6-well plates and infected at a MOI of 0.01 TCID_50_/cell as described previously(41). Infectivity titres in the samples were then determined by TCID_50_ assay.

### Ethics statement

Ferrets (weight range 600–1500 g) were housed in the Bioresources Facility at the Peter Doherty Institute for Infection. Experiments were conducted with approval from the University of Melbourne Biochemistry & Molecular Biology, Dental Science, Medicine, Microbiology & Immunology, and Surgery Animal Ethics Committee, in accordance with the NHMRC Australian code of practice for the care and use of animals for scientific purposes (project license number 1313040).

### Ferret experiments

Male and female ferrets aged 6-12 months old were used in experiments. All animals received food and water ad libitum. All test animals were confirmed to be seronegative to circulating human influenza strains using a hemagglutination inhibition assay prior to the experiments. The replication and contact transmission efficiency of the wild-type and variant viruses in the ferret model was tested by experimental infection of ferrets with a ‘pure’ population of wild-type (WT) or mutant (MUT) viruses as described below. In independent experiments, ferrets were infected with ‘mixtures’ of wild-type and mutant viruses to determine the relative fitness of each in ‘competitive mixtures’ experiments (41, 45, 46). Viral stocks were standardized to 1×10^5^ TCID_50_/ml in Dulbecco’s Phosphate Buffered Saline (Sigma, Australia) for ferret inoculation. To make viral mixtures for WT-H273: MUT-Y273 and WT-D197: MUT-N197, the standardized stocks were volumetrically mixed at 80:20, 50:50 and 20:80 ratios.

On day 0, donor ferrets were anaesthetized by intramuscular injection with 20 mg/ml Ilium Xylazil-20 (Troy-laboratories, Australia) and intranasally inoculated with 0.5 mL of (i) pure populations of WT (n=4) and MUT (n=4) viruses, or (ii) mixtures at ratios of 50:50 (n=4), 80:20 (n=2) and 20:80 (n=2) of WT and MUT viruses. The ferrets were then housed in high-efficiency particulate air (HEPA) filtered cages. After 24 hours (Day 1 p.i.), one naïve recipient ferret was co-housed with one donor ferret to allow for contact transmission. The donor and recipients were co-housed for the entire length of the experiment (11 days). Nasal washes were collected from donors and recipients each day throughout the experiment and temperature and weight was monitored daily, as described previously (45). All ferrets were euthanized on Day 11. Infectious viral load in ferret nasal washes were quantified by TCID_50_ assay.

### RNA extraction and quantitative analysis of viral RNA in ferret nasal washes

The viral RNA from ferret nasal wash was extracted and quantified using real-time PCR. The real-time primers were provided by the CDC (Atlanta, USA), and capable of detecting the influenza B NS gene. Internal controls with known RNA copy numbers of B/England/598/2014 NS transcripts were generated in-house and included for each assay. This was used to correlate cycle threshold (Ct) with RNA copy number and quantitate viral RNA in nasal wash samples.

### Molecular analysis of ferret nasal washes

The proportion of WT and MUT viruses in each nasal wash sample was determined by pyrosequencing (45) using primer pairs specifically designed for PCR amplification and sequencing analysis (Table S3). To validate the performance of the assay, pure populations and known mixtures for each WT: MUT pair were serially diluted and tested across three separate assays. Based on these validation tests, only samples with greater than 4.3 Log_10_ NS gene copies/µL were analysed for mixture estimation.

Next generation sequencing was completed on nasal wash samples from the last day of viral shedding for each recipient in the pure population study. If the viruses did not transmit, then nasal wash samples from the last day of shedding of donors were analysed. This was done to confirm genetic stability of the viruses after replication and transmission. In competitive mixture experiments, where pyrosequencing results indicated a large increase in MUT viruses in recipient ferrets, nasal wash samples from the last day of viral shedding was also analysed.

RT-PCR amplification of all influenza B genes was done on extracted viral RNA using universal influenza B primers and PCR conditions as described in Zhou et al 2014 (47). Illumina library preparation and sequencing (Miseq v2 with 150bp paried end-read) was done off-site at the Murdoch Children’s Research Institute, Melbourne, Australia. The NGS reads were mapped to their respective WT/MUT genome using Bowtie2 v2.2.5(- very-sensitive-local) (http://bowtiebio.sourceforge.net/index.shtml) after building an index of the references with the bowtie2-build program. SAM tools v1.7 (https://sourceforge.net/projects/samtools/files/samtools/1.7) was used to process sequence alignments and generate pileup files. The pileup files were then used to build consensus sequence of all genes and scanned for minorities using QUASR v6.08 (https://sourceforge.net/projects/quasr/) (48).

### Quantitative assessment of differences in virus replication and transmission fitness

The fitness of a virus depends on its ability replicate within a host and to transmit between hosts. We used a mechanistic model of viral replication and competition to assess the relative within-host replicative fitness of the variant compared to the wild-type virus (45, 49).

As previously described, our model expanded on the standard Target Cell - Infectious Cell – Virus (TIV) model of viral dynamics (49). The viral population is stratified by strain and accounts for infectious and non-infectious viral material by simultaneously fitting to TCID_50_, real-time PCR and pyrosequencing data (50).

From the model fit, we estimate the ratio of the basic reproduction number, *R*_*0*_, of the two viral strains in the competitive mixture, where *R*_*0*_ describes the expected number of secondary infections produced by a single infected cell in a population of susceptible cells (51).

To assess the relative ability of the variant to transmit between hosts we fitted a transmission model as described in McCaw et al. to pyrosequencing data (46). We utilized the pyrosequencing measurements of the proportion of mutant virus in the donors (on the inferred day of infection) and the recipients. We extended on our previous work to account for pyrosequencing measurement errors in producing our estimates. The transmission model has a single parameter describing the relative transmission fitness of the two strains where a value greater than 1 indicates enhanced transmissibility of the mutant.

### Statistical analysis

Viral titres between different groups from the *in vitro* and *in vivo* experiments were analysed by a two-way ANOVA with Bonferroni’s post-hoc analysis using GraphPad Prism 5.0. All other comparisons such as Area under the Curve (AUC), peak viral titres, NA activity and K_m_ values were made using non-parametric Mann Whitney’s U test. Parameters for the mathematical models used to determine relative finesses (both within and between hosts) were estimated from the data in a Bayesian framework using the R interface to STAN (52, 53) (see Text S4).

## Acknowledgements

The Melbourne WHO Collaborating Centre for Reference and Research on Influenza is Supported by the Australian Government Department of Health

We gratefully acknowledge GISRS laboratories for providing influenza viruses to the Melbourne WHO Collaborating Centre and Thomas Cumming, Celeste Tai, Leonard Izzard, Ding Yuan Oh, Rebecca Jayne Bower and Charlene Plascenia for assistance with animal experiments.

